# Znf804a is a regulator of circadian behaviors in zebrafish

**DOI:** 10.64898/2026.04.29.721668

**Authors:** Brandon L. Bastien, Eric H. Li, Mary E. S. Capps, Summer B. Thyme

## Abstract

Sleep disturbances are common among individuals with schizophrenia and can exacerbate disruptions in cognitive processes like learning and memory. Elucidating pharmacologically targetable molecular pathways perturbed by schizophrenia genes may uncover new treatment avenues. Here, we investigated the relationship of the schizophrenia-associated gene *znf804a* with sleep and circadian pathways. Using multi-day behavior tracking, we showed that *znf804a* zebrafish mutants displayed changes in sleep and circadian behaviors when light cues were removed. Through bulk RNA sequencing of fish raised under normal light cycling and dark-only conditions, we identified altered gene expression in the core and auxiliary pathways controlling circadian rhythms. Expression of *fbxl3a*, which encodes a modulator of the core negative feedback regulator of the clock, decreased in a dose-dependent manner as *znf804a* mutant copy number increased. Further analysis also revealed shifts in the relative abundance of specific transcripts, including *idh1*, suggesting *znf804a* could influence transcript processing or stability. Together, these findings link a *ZNF804A* ortholog to sleep and circadian behaviors and identify the regulation of *fbxl3a* and transcript processing as candidate mechanisms through which this schizophrenia risk gene may influence circadian biology.

## Introduction

Neuropsychiatric conditions, such as schizophrenia (SCZ), are a leading cause of disability, with SCZ alone affecting 1% of individuals worldwide^1^. Sleep disturbances is a common symptom in individuals with SCZ and can cause major disruptions in everyday life^2^ by affecting memory consolidation, attention, decision-making, and other cognitive processes^3,4^. SCZ is highly heritable, and identifying how SCZ-associated genes influence sleep may reveal molecular pathways that can be pharmacologically targeted to improve sleep and cognitive function.

*ZNF804A* is among the top genes associated with SCZ, first identified in a genome-wide association study (GWAS)^5^, and further replicated in other cohort studies^6–9^. Several human studies have also linked a risk allele (*rs1344706*) in the second intron of *ZNF804A* to brain anatomy^10–12^ and connectivity^13–15^. The *ZNF804A* risk allele also has substantial effects on learning and memory. One study found that individuals with SCZ carrying the risk allele of *ZNF804A* performed worse on a visual memory task than those without^16^. Notably, the *ZNF804A* risk allele has been linked to changes in sleep neurophysiology and impacts the relationship between sleep and memory consolidation^17^. Taken together, these findings suggest that *ZNF804A* influences neural circuitry and sleep-dependent neural processes in individuals with SCZ.

Several *in vitro* and animal model studies have demonstrated that *ZNF804A* and its orthologs have roles in cellular and molecular processes in the nervous system. At the circuit level, ZNF804A regulates neurodevelopment and plasticity, including by altering the expression of genes linked to cell adhesion and neural migration^18^ and recruiting glutamatergic receptors^19^ and translation machinery to synapses^20,21^. However, its specific molecular and biochemical functions remain unclear. Although it is predicted to contain a C2H2-type zinc finger domain commonly found in transcription factors^22^, and one study found it binds to DNA to regulate transcription^23^, it has more broadly been implicated in mRNA splicing and transcript processing. For example, ZNF804A has been shown to regulate mitochondrial genes and genes regulating translation^20^, and it interacts with RBFOX1, a clinically relevant mRNA splicing factor^24^. These findings point towards ZNF804A having roles in RNA processing and translation to coordinate neural development and circuit plasticity.

Links between sleep and *ZNF804A* in model organisms have not yet been established; however, effects on brain function and behavior have been reported. Mutant mice displayed impaired learning and memory in behavioral tasks and increased anxiety and depressive-like behaviors^25,26^, along with synaptic plasticity deficits in the hippocampus and glutamate/GABA imbalances^25^. One study also found sex-dependent differences in hippocampal LTP and neuromodulation, including serotonin and dopamine signaling^26^, in mutants. In zebrafish, *znf804a* regulates brain activity, with midbrain hypoactivity reported in *znf804a* mutants compared to siblings^27^, although no strong behavioral differences were observed under normal light-cycling conditions in larvae. Orthologs of *ZNF804A* therefore play important roles in the brain and behavior in humans and model organisms. Because sleep can affect anxiety-related behaviors and learning and memory, linking these *ZNF804A* orthologs to sleep may provide insights into other phenotypes observed in model organisms.

In this study, we aimed to understand the mechanisms underlying the *znf804a* mutant brain activity phenotypes^27^ and how *ZNF804A* contributes to psychiatric disorders. Through a combination of behavioral and transcriptomic studies, we determined that loss of *znf804a* disrupts molecular pathways controlling circadian rhythms, leading to altered circadian behaviors.

## Results

### *znf804a* mutants display disrupted circadian activity

Given the relationship between *ZNF804A* and sleep in humans, we were interested in investigating circadian behaviors and pathways in *znf804a* zebrafish mutants. Preliminary bulk RNA sequencing (**Figure S1**) suggested that loss of *znf804a* reduced the expression of the circadian genes *fbxl3a* and *per1b*, which inspired us to further test circadian behaviors. We monitored 4 dpf larvae in our 96-well behavior assay under four conditions: normal 14 hr light/10 hr dark cycles^27^, dark/dark cycles beginning at 10 PM (the start of their normal dark period), light/light cycles, and a light/dark experiment, hereafter referred to as light shift, where their light period was shifted to start at 3 AM and end at 5 PM after normal circadian entrainment. In normal light/dark cycles, we saw that there was a small shift in circadian activity in *znf804a* mutants compared to wild-type siblings (**Figure 1A**). We repeated this experiment under dark/dark conditions, and observed a stronger shift in activity and decreases in overall activity (**Figure 1B**). These results were replicated in two independent clutches of fish five generations later (**Figure S2A**). We then raised fish under constant light conditions, and found that, while there was not a circadian drift, there was a reduction in activity, particularly on the second day of recording (**Figure 1C**). In the light shift experiment, we saw that shifting the circadian cues caused the wild-type siblings to compensatory shift their activity, whereas the mutants were delayed in compensating for their waking activity and sleep bouts (**Figure 1D**). There was no strong heterozygous circadian phenotype, although the heterozygous fish did have lower activity in four of the six behavioral runs (**Figure S2B**). Overall, these experiments demonstrate that *znf804a* mutants exhibit changes in sleep and circadian phenotypes, which emerge most strongly when fish are deprived of light.

**Figure 1:**
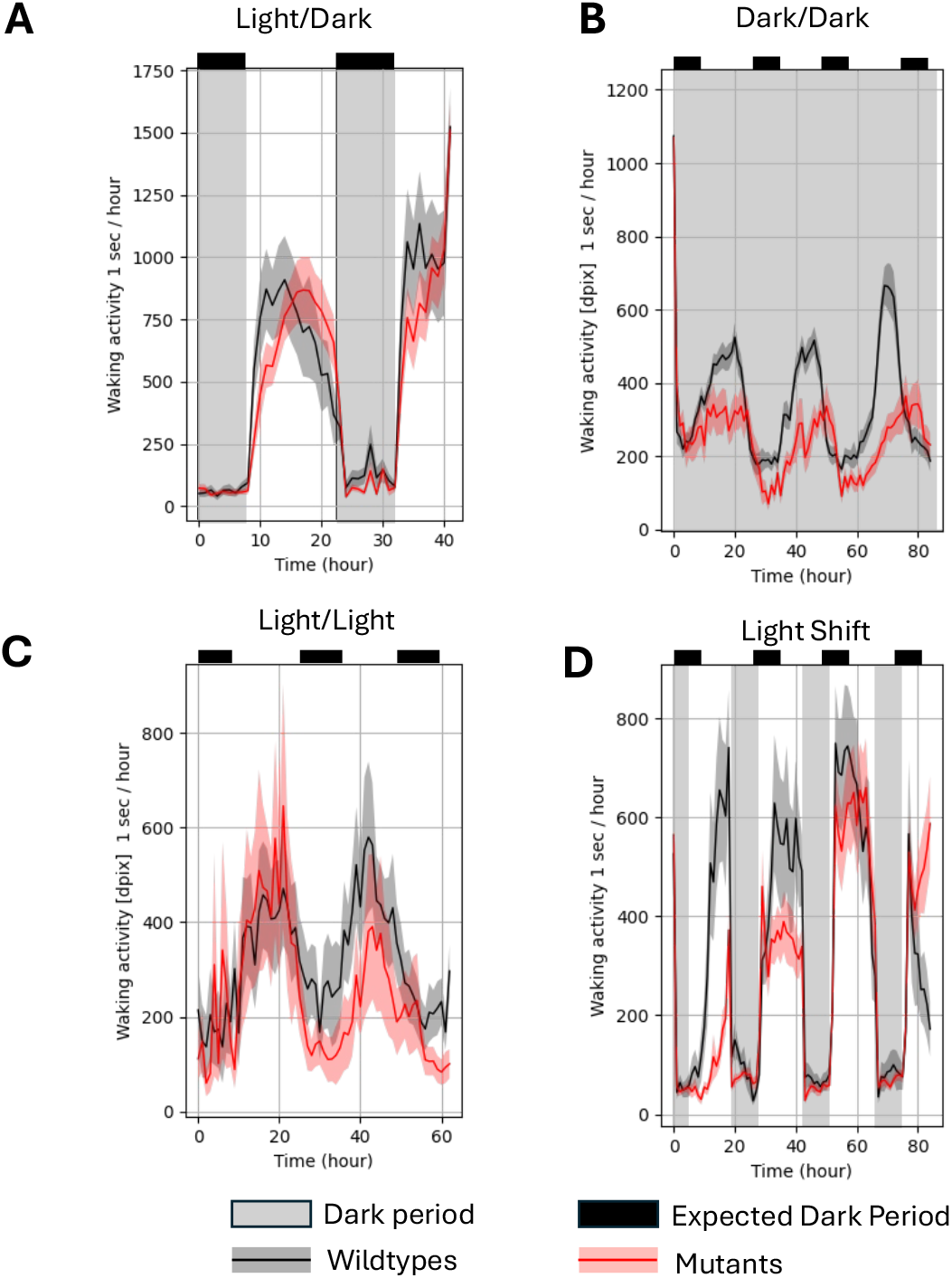
*znf804a* mutants exhibit circadian behavior disruptions. **A**) wildtype siblings (black) (n=15) and *znf804a* mutants (red) (n=22) waking activity binning by hour over the course of the light/dark experiment starting at 4 dpf, in which normal light/dark cycles were maintained. The mutants exhibit a small but noticeable shift in their activity during the first 24 hours of this experiment. **B**) binned wildtype (n=25) and mutant (n=12) waking activity in the dark/dark experiment, where the mutants display reduced waking activity and circadian drift that gets larger with each day of the experiment. **C**) binned wildtype (n=16) and mutant (n=10) waking activity in the light/light experiment, where circadian drift is not observed, but reduced waking activity is seen in the mutants, particularly on the second day of the experiment. **D**) binned wildtype (n=9) and mutant (n=20) waking activity in the light shift experiment, where wild-type fish can maintain activity during light periods while mutants exhibit a shift in activity during the first day of the experiment, reduced activity during the second day, before nearing control levels of activity during the third day.

### Circadian genes are dysregulated in *znf804a* mutants

To determine the molecular effects in *znf804a* mutants that underly the observed behavioral phenotypes, we raised fish either in normal light/dark cycle conditions or in the dark starting at 2 dpf and performed RNA sequencing on larval heads at 6 dpf. We compared mutants and wildtypes within each condition, as well as within-genotype differences across conditions to generate a list of differentially expressed genes (DEGs). Using strict thresholding (adjusted p-value <0.05), we identified 86 DEGs between mutants and wildtypes when raised in normal light/dark conditions (**Table S1**) and 184 DEGs when raised in total darkness (**Table S2**). Development in the dark also altered gene expression in both wildtypes and mutants, with wildtypes having 685 DEGs (**Table S3**) and mutants having 2512 DEGs in the dark compared to the light (**Table S4**) (**Figure 2A**). These results indicate that both *znf804a* and rearing in a light/dark cycle have substantial effects on gene expression.

**Figure 2:**
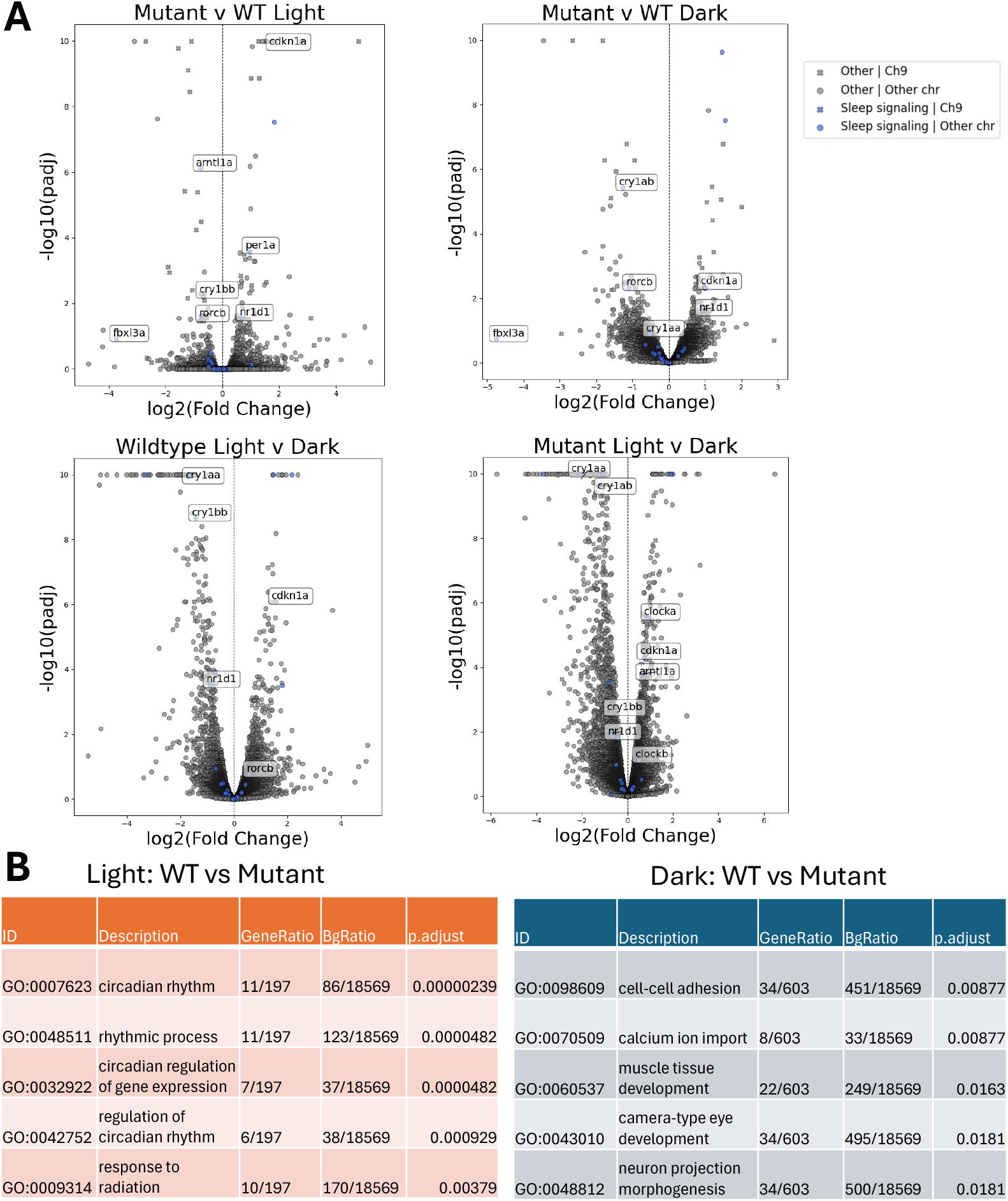
Gene expression changes in *znf804a* mutants and light conditions. **A**) Volcano plots depicting differentially expressed genes for wildtypes vs mutants raised in normal light/dark conditions (top left) or dark/dark conditions (top right) or differences between fish raised in light/dark and dark/dark conditions for wildtypes (bottom left) and mutants (bottom right). Each condition consists of 4 samples, which were made through pooling 4 larval heads. **B**) Table depicting top terms identified in gene ontogeny (GO) analysis.

Unbiased Gene Ontology (GO) enrichment analysis revealed pathways that were impacted by the loss of function of *znf804a* and rearing in normal light/dark conditions compared to the dark. Consistent with our initial findings (**Figure S1**), circadian rhythm and related terms were enriched in DEGs between mutants and wiltypes raised in a regular light/dark cycle. These circadian terms were not detected in DEGs between mutant and wild-type fish raised in the dark. However, several GO terms related to neural development, cell adhesion, and muscle development were identified, suggesting *znf804a* plays direct or indirect roles in development (**Figure 2B**). Notably, the mutant fish were less active than wildtypes in behavior experiments when raised in the dark (**Figure 1B, Figure S2**), so some transcriptomic differences could also be a downstream consequence of inactivity. When comparing each genotype across light/dark conditions, both genotypes exhibited broader alterations in gene expression. The wild-type fish had several expected differences in gene categories, including those related to circadian rhythms and responses to external stimuli. Mutants also had gene expression changes in circadian and environmental stimuli categories; however, several muscular and neurodevelopmental terms not seen in the wildtypes were also significant in the mutants (**Table S5**). Changes in activity in regular light/dark conditions between mutants and wildtypes were not seen previously^27^, so this could again be a consequence of reduced activity in the dark.

The core molecular mechanisms controlling circadian rhythms consist of positive regulators (*clock* and *BMAL*/*arntl* genes) and negative regulators (*per* and *cry* genes). Additional mechanisms that modify this pathway include the REV-ERB pathway (*nr1d* genes), retinoic orphan response genes (*ror* genes), and *fbxl3* genes (**Figure 3A, Table S6**)^28–31^. We next compared how *znf804a* mutations and exposure to light/dark impacted each of these elements from the RNA sequencing data. We found that genes from each of these molecular pathways were impacted in *znf804a* mutants (**Figure 3B**). In the core pathway, *arntl1a, cry1bb*, and *per1a* were dysregulated in *znf804a* mutants. Additionally, *clocka, clockb*, and *arntl1a* were significantly higher in mutant fish raised in the dark vs those compared in normal light conditions. Several genes were affected by light exposure, including *cry1aa, cry1bb, per2*, and *per3*, consistent with prior reports^32,33^. These genes may allow the *znf804a* mutants to maintain circadian rhythms in normal light/dark conditions (**Figure 1A**). In the REV-ERB pathway, *nr1d1, nr1d4a*, and *nr1d4b* were dysregulated by both *znf804a* mutation and by exposure to light (**Figure 3B, Table S6**). In the retinoic acid orphan response group of genes, only *rorcb* was disrupted in *znf804a* mutants. Light/dark conditions did not affect the expression of any of the *rora, rorb*, or *rorc* genes. We next compared the expression of *cdkn1a*, an E-box-rich gene influenced by circadian genes like the *clock* and *arntl*, across all four groups, and found its expression was dysregulated by both *znf804a* mutations and light/dark conditions. Thus, *znf804a* mutations appear to influence RNA expression of genes related to the canonical and auxiliary circadian pathways, which could explain the circadian phenotypes seen in the fish.

**Figure 3:**
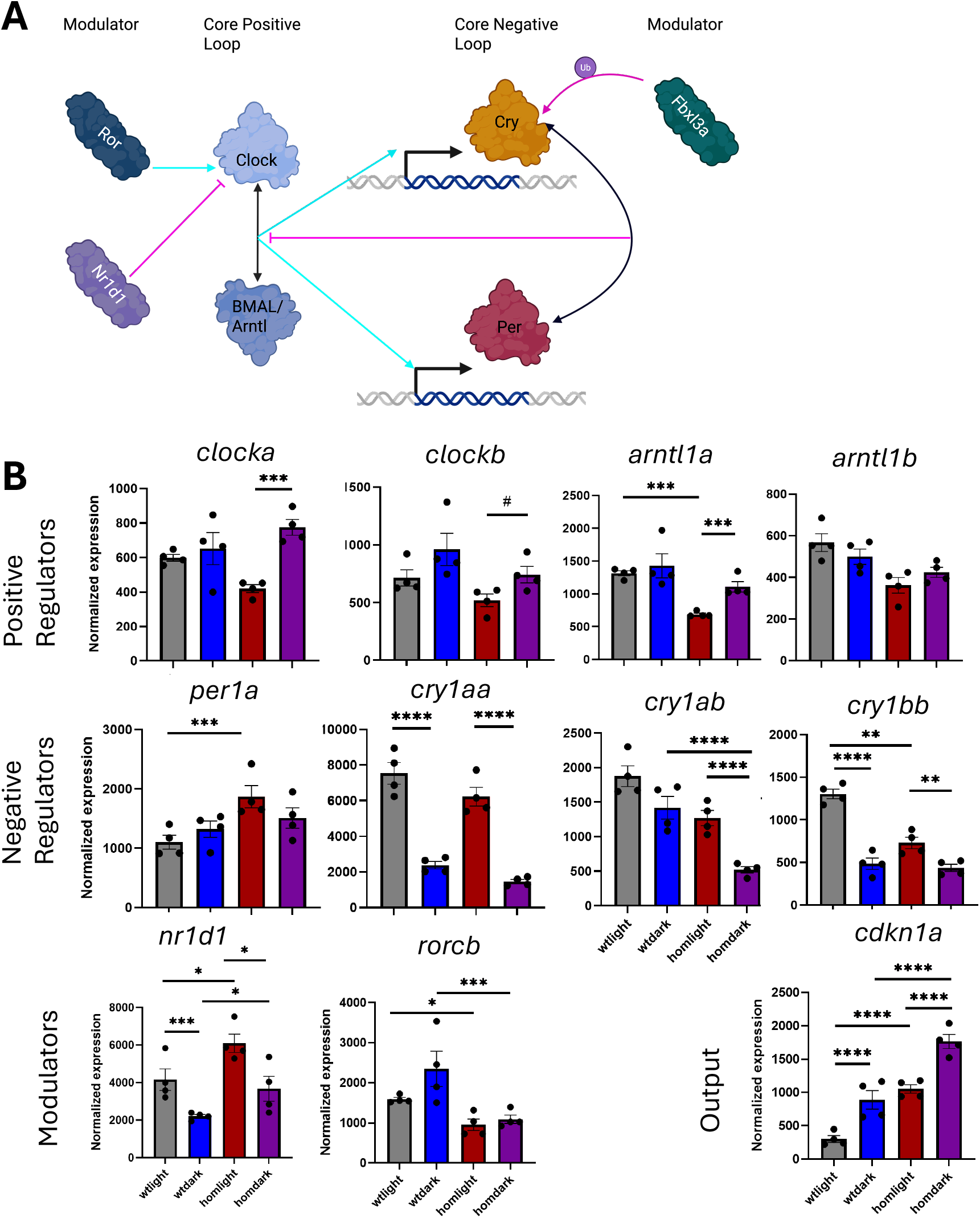
Circadian gene expression alterations across genotypes and light conditions. **A**) Diagram of circadian genes. The core molecular pathway controlling circadian rhythms consists of the *CLOCK* and *BMAL* genes acting together through forming heterodimers and activating transcription, while period/*PER* and *CRY* act as repressors by disrupting *CLOCK* and *BMAL*. An additional pathway, the REV-ERB pathway, which comprises *NR1D* genes, also represses *CLOCK* and *BMAL* function. Additional auxiliary modulators include the retinoic acid orphan receptors (*ROR* genes), which interact with the REV-ERB pathway ^29^, and *FBXL* genes, which can influence degradation of the CRY genes^30^. **B**) Normalized gene expression of select circadian genes for mutants and wildtypes in normal light/dark or dark/dark conditions. Significance denoted is the result of correcting for multiple comparisons after differentially expressed gene analysis: #=p<0.1, ^*^p<0.05, ^**^p<0.01,^***^p<0.001, ^****^p<0.0001.

### *fbxl3a* has a gene-dosage relationship with *znf804a*

One notable circadian gene whose expression was almost eliminated in *znf804a* mutants was *fbxl3a* (**Figure 4A**), which encodes a protein that regulates CRY stability^30^. A previous zebrafish study found that loss of *fbxl3a* resulted in circadian drifting when larvae were removed from normal light/dark cycles^31^, similar to our observed phenotype (**Figure 1B**). Heterozygous fish exhibited an intermediate reduction in *fbxl3a* expression compared to the wild-type and homozygous mutant siblings. To validate this relationship, we injected wild-type embryos with two cocktails of guide RNAs and Cas9 mRNA targeting the protein-coding or regulatory region of *znf804a*. We found significant or trend-level reductions in *fbxl3a* expression in these two independent crispant experiments (**Figure S3**). *Nr1d1* expression was also increased, confirming broad effects on circadian gene regulation when *znf804a* levels are reduced. We observed similar patterns for many of the individual *fbxl3a* transcripts (**Figure 4B**). Based on a comprehensively annotated transcriptome from the Lawson Lab^34^, the large TCONS_00125506 isoform and the larger of the protein-coding containing isoforms, TCONS_00125508, both exhibited a similar gene-dosage dependent reduction in expression across wild-type, heterozygous, and mutant fish as overall *fbxl3a* expression. This supports the global reduction in *fbxl3a* gene expression seen in the gene-level analysis. Transcript TCONS_00125510, the isoform containing only the protein coding region and no UTRs, was downregulated in the mutants and heterozygous fish compared to the wildtypes, which does seem to indicate that *znf804a* mutants and heterozygous fish have deficiencies in processing this short transcript. Transcript TCONS_00125507 and 09, which are lowly expressed, did not differ between *znf804a* mutants, heterozygous, and wild-type siblings (**Figure 4C**). These results suggest that overall *fbxl3a* expression is reduced in a gene-dosage-dependent manner by *znf804a* loss, and that even in heterozygous fish, transcript processing of *fbxl3a* may be altered.

**Figure 4:**
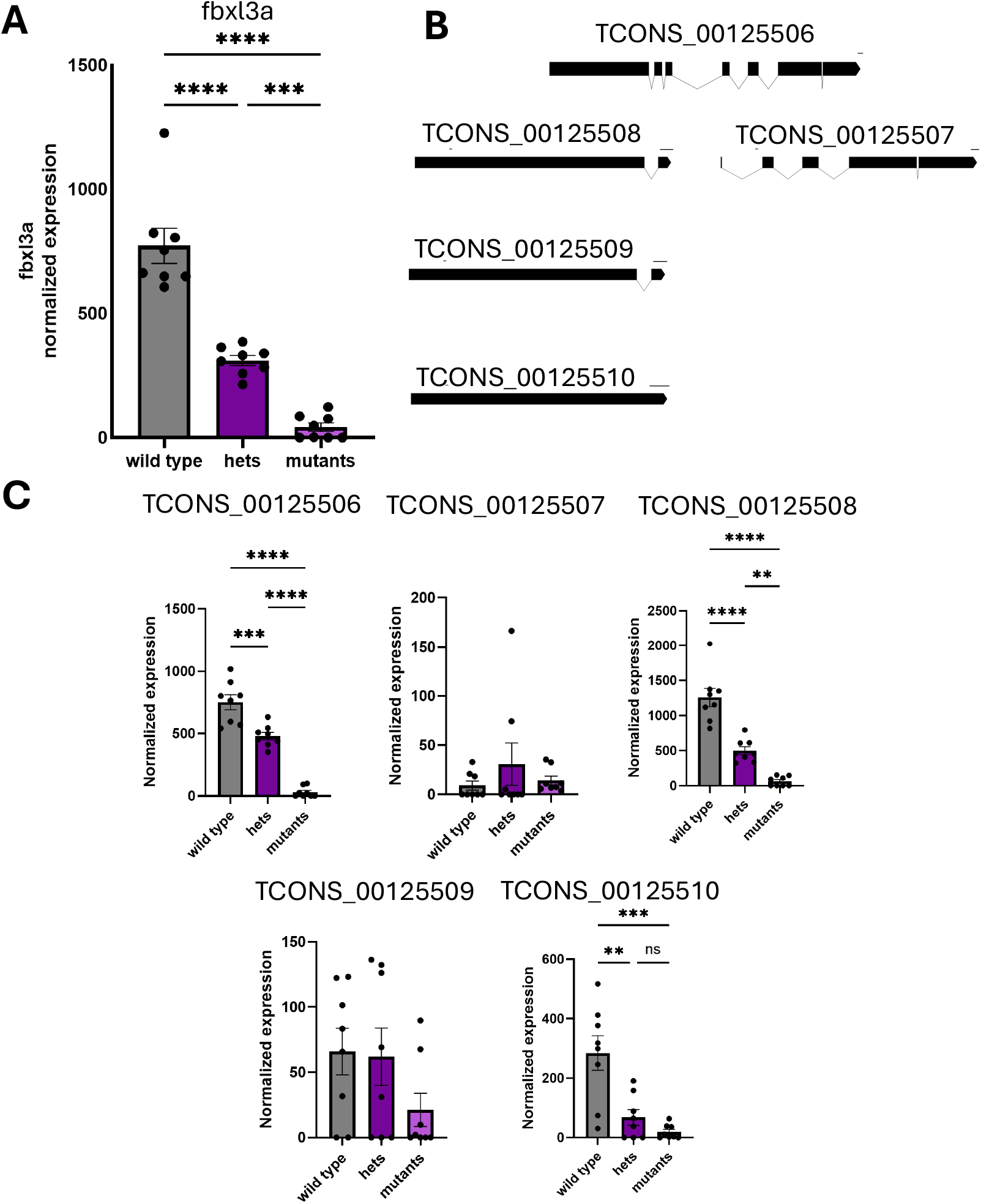
*fbxl3a* expression is influenced by gene dosage of *znf804a*. **A**) Normalized expression of *fbxl3a* in pooled mutant, wild-type, and heterozygous fish across all lighting conditions. In light and dark conditions separately, there were reductions in expression from wildtype to heterozygous (-1.17 log_2_ fold and -1.58 log_2_ fold in light and dark, respectively) and heterozygous to mutants (-2.55 log_2_ fold change and -3.19 log_2_ fold change in light and dark respectively) that were narrowly not significant after multiple corrections, but p-values for pooled comparisons were <0.001 and the points separate into three distinct groups. **B**) Diagram of annotated transcripts of *fbxl3a* identified in the RNA sequencing data, with exons depicted as rectangles and introns depicted as diagonal lines. Based on the similarity of the transcripts, TCONS_00125507 (3118 bps) and TCONS_00125508 (1947 bps) are spliced from TCONS_00125506 (5881 bps), with TCONS_00125510 (1297 bps) and TCONS_001255109 (1831 bps) being further splice isoforms derived from TCONS_00125508. TCONS_00125508-10 has the protein-coding region of the gene and differs in 5′ and 3′ UTRs. TCONS_00125507 likely seems to be an intermediate.**C**) Normalized expression of all of the transcripts of *fbxl3a* in pooled mutant, wild-type, and heterozygous fish across all lighting conditions. Results of One-Way ANOVA: ^*^p<0.05, ^**^p<0.01, ^***^p<0.001, ^****^p<0.0001.

### Transcript abundance shifts are observed in several genes, including genes linked to sleep and circadian pathway modulation

While the exact function of *znf804a* is unknown, *znf804a* has been highly implicated in RNA processing^20,24^. Further analysis of individual transcripts in our RNA sequencing data revealed multiple transcripts within a gene, in which one transcript was upregulated while another was downregulated (**Table S7**). These shifts may reveal differences in transcript processing. In the fish raised in normal light conditions, 12 genes showed shifts in transcript abundance in *znf804a* mutants. Most of these pairs of isoforms had minor differences, generally less than 100-200 bps differences in total length, with several of the longer sequences having 5′ modifications. However, 4 of the genes – *ctnnd2b, caska, RAMP1*, and *idh1* – exhibited shifts involving substantially larger differences in isoforms. Notably, in *idh1*, the long transcript *idh1*-201/ TCONS_00123632 was not detected in any of the wild-type fish, but was present in all mutant samples and 3 of the 4 heterozygous samples. Correspondingly, an intermediate transcript of 3490 base pairs not annotated on ENSEMBL was elevated in the wildtypes compared to the mutants. The smaller *idh1*-202 transcript was unaffected (**Figure 5A-C**). *caska* and *ctnnd2b* also exhibited similar shifts in expression of one transcript to another in the mutants compared to wildtypes (**Figure 5D**). These results suggest that *znf804a* may influence transcript processing or stability.

**Figure 5:**
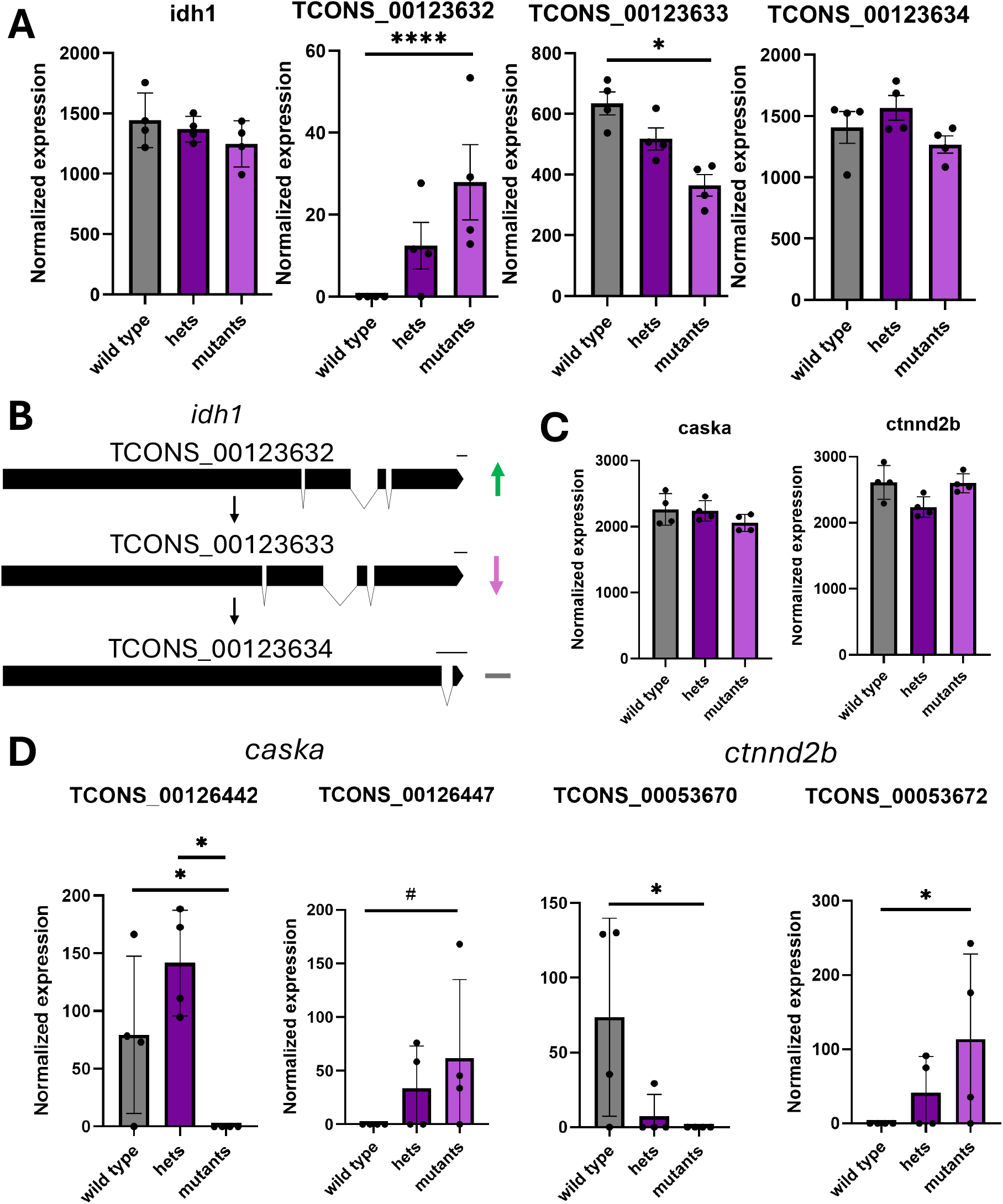
Transcript-specific shifts in gene expression of sleep and circadian-related genes. **A**) Normalized expression of *idh1* (far left) and its three annotated transcripts (right). **B**) Diagram of all three annotated *idh1* transcripts, with exons depicted as rectangles and introns depicted as diagonal lines. Normalized gene expression of **C**) overall *caska* and *ctnnd2b* gene expression and **D**) select annotated transcripts of *caska* and *ctnnd2b* with opposing shifts in expression. Significance denoted is the result of correcting for multiple comparisons after differentially expressed transcript analysis: ^*^p<0.05, ^**^p<0.01, ^***^p<0.001, ^****^p<0.0001.

## Discussion

In this study, for the first time in model organisms to our knowledge, we demonstrate a link between the SCZ risk gene ortholog *znf804a* and circadian rhythms. Behaviorally, zebrafish with mutations in *znf804a* exhibited disrupted circadian behaviors when light cues were removed. At the molecular level, *znf804a* mutants exhibited disruptions in circadian pathways both under normal lighting conditions and in the dark. These changes were widespread across all of the core and auxiliary circadian pathways. These two lines of evidence align with findings that *ZNF804A* impacts sleep in humans^17^. While uncertainties regarding ZNF804A’s exact function remain due to its ambiguous protein structure and biomolecular interactions, we suggest two hypotheses for how *znf804a* impacts circadian pathways:

One hypothesis for how *znf804a* might influence circadian regulation is that Znf804a is directly acting on *fbxl3a*, which encodes an auxiliary regulator of circadian rhythms. The expression of *fbxl3a* is massively reduced in a gene-dosage-dependent manner, and analysis of individual transcripts revealed that several followed this pattern (**Figure 4**). The loss of *fbxl3a* without upregulation of any annotated *fbxl3a* transcripts suggests that *fbxl3a* is either not being transcribed or precursor transcripts are not being processed and are eliminated via nonsense-mediated decay. The loss of *fbxl3a* would reduce the regulation of CRY through ubiquitination, which could cause a shift in cycling of circadian genes and shift circadian activity, which is seen in both *znf804a* and *fbxl3a* mutants in the absence of circadian cues^27^.

A second, non-mutually exclusive hypothesis is that *znf804a* indirectly influences circadian genes either through direct transcript processing of other genes or by impacting the development or function of neurons in circadian circuits. There were four genes in which two transcripts of a single gene with substantial differences, ranging from hundreds to thousands of base pairs, were shifted in opposing directions. Although speculative, this and similar transcript-level changes could contribute to disruptions of circadian pathways and genes, especially given the links between *CASK* and *CTNND2* in sleep regulation^35,36^ and *IDH1*’s regulation of circadian pathways through Smad signalling^37^. While it is difficult to infer whether Znf804a is directly interacting with these gene products, the shift in transcripts seen in these genes, coupled with *znf804a*’s proposed roles in RNA splicing, suggests that *znf804a* could be responsible for these imbalances. These transcript imbalances could have downstream effects on circadian pathways and genes, such as on *fbxl3a*, and further support that Znf804a could also be directly regulating *fbxl3a* transcripts. Finally, although we cannot rule out the possibility of developmental circuit effects in homozygous mutants, we suggest that this cause is unlikely because 1) there are no morphological changes observed in the mutant brain^27^, 2) the transcriptomics did not indicate any aberrant neuron types through GSEA analysis^38^ (**Figure S4, Table S8**), 3) *znf804a* is broadly expressed throughout the entire brain, making it less likely to affect only specific cell types^39^, and 4) isoform-level differences were observed in the heterozygous compared to the wild-type, even in the absence of strong heterozygous transcriptomic changes (**Tables S9-S12**).

Additional studies elucidating how the exact functions of *znf804a* will provide further clarity of how *znf804a* impacts circadian genes. There is currently not a resolved protein structure or AlphFold model^40^, which makes it unclear whether *znf804a* is directly binding to RNA with its zinc finger domain or acts as a scaffold. To establish more causal relationships between *znf804a* and the differential gene expression seen here, approaches like RNA base editors^41^, could be used to narrow which transcripts Znf804a interacts with. Alternatively, TurboID proximity labeling^42^ could be used to identify interacting partners if *znf804a* acts as a scaffold could be used to narrow protein partners. Linking misregulation of these transcripts to disruption of circadian pathways will help narrow molecular pathways that could be targeted to ameliorate phenotypes like sleep deficits.

The prevalence of sleep disturbances in individuals with SCZ and the interconnectedness of sleep and cognitive behaviors like learning and memory necessitate a better understanding of how SCZ risk genes modulate sleep and circadian rhythms. Our results highlight the utility of model organisms to bridge SCZ genes and perturbations in sleep-related molecular pathways/behaviors, which generated testable hypotheses and identified pathways that can be prioritized in drug discovery. Further studies in model organisms can examine how *ZNF804A* ortholog mutations disrupt sleep architecture to determine if *ZNF804A* has conserved roles in sleep-related neurophysiological processes. Further, assessing how sleep disturbances in *Znf804a/znf804a* mutants affect learning and memory could further bridge findings in model organisms to findings in humans.

## Materials and Methods

### Zebrafish Husbandry

Zebrafish experiments were approved by the UMass Chan Institutional Animal Care and Use Committee (IACUC protocols IPROTO202300000053). All animals were maintained on a 14 h/10 h light/dark cycle at 28°C before experiments. The process for generating the mutants used in this study is described in Thyme et al. 2019, and they were of the EK background. Larval fish were grown in 100 mm petri dishes at a density of 50 fish/dish. Debris was removed before experiments, and only healthy fish with swim bladders were included in this study.

### Larval Behavior

Larval behavior assays were performed and analyzed as described^43^, with stimulus response paradigms removed. An IR lamp emitting 960 nm wavelength light (past the visible spectrum for zebrafish) and a Grasshopper GigE camera with 50 mm fixed focal length lens and an IR filter was used for tracking, and the light level was controlled through a white LED panel. For the larval experiments, the LED panel was controlled from a command file and was either turned off at 10 PM and turned on at 8 AM (14 hr light/10 hr dark), turned off at 10 PM and remained off for the duration of the experiment (Dark condition), remained on for the duration of the experiment (Light condition), or was shifted to turn on at 3 AM and turn off at 5 PM. Tracking was performed using LabVIEW. Behavioral analysis code is available at: https://github.com/thymelab/ZebrafishBehavior.

### RNA Sequencing and Analysis

RNA sequencing was performed as previously described^38^. Zebrafish were raised either under normal light/dark conditions or in the dark starting at 2 dpf in 12 well plates at a density of 12 fish/well. Heads were collected from anesthetized larval zebrafish (Syncaine MS 222 Fish Anesthetic) and kept at -80°C until sample preparation. Bodies were collected for PCR genotyping. RNA samples consisting of 4 pooled fish were combined and extracted for a total of 4 samples/condition (wild-type, hets, and mutants genotypes in light/dark and dark conditions).

RNA extractions were performed using E.Z.N.A. MicroElute Total RNA Kit (Omega Bio-Tek R6834-02). Samples were nanodropped on a ThermoScientific Nanodrop One Spectrophotometer to determine RNA concentration and purity.

Reverse transcription was performed by using 6 uL of RNA sample with 0.3 μL of 10 μM RT Oligo (5′-AGACGTGTGCTCTTCCGATCT(30)VN-3′), 3 μL of 10 mM dNTP mix (Thermo-Fischer, R0192), 0.3 μL of an RNase Inhibitor 40 U/μL (Life Technologies, AM2694), and 2.5 μL of 1 M Trehalose (Life Sciences, TS1M-100). Samples were incubated at 72°C for 3 minutes and then placed on ice. 6 μL of 5X Maxima RT Buffer (Thermo-Fischer, EP0751), 0.3 μL of Maxima RNase H-Minus RT 200 U/μL (Thermo-Fischer, EP0751), 10.35 μL of 1 M Trehalose (Life Sciences TS1M-100), 0.3 μL of 1 M MgCl2 (Invitrogen, AM9530G), 0.3 μL of 10 μM TSO (5′-AGACGTGTGCTCTTCCGATCTNNNNNrGrGrG-3′), and 0.75 μL of RNase Inhibitor 40 U/μL (Life Technologies, AM2694) was added, gently mixed with a pipeptte, and incubated for 90 minutes at 50°C and 5 minutes at 85°C to heat inactivate. Finally, 7 μL of this reaction was added to 5 μL of nuclease-free water, 0.5 μL of 10 μM PCR Oligo (5′-AGACGTGTGCTCTTCCGATCT-3′), and 12.5 μL of Kapa HiFi HotStart PCR ReadyMix (KAPA Biosystems, KK2601). A whole transcriptome amplification PCR was performed using one cycle of 98°C for 3 minutes, 14 cycles of 98 °C for 15 seconds, 67°C for 20 seconds, and 72°C for 6 minutes, and a final extension of 72°C for 5 min. Products were then purified using a 0.8X AMPure XP SPRI beads (Beckman-Coulter, A63881). DNA was eluted in 10 μL elution buffer, and sample concentration was determined with the Promega QuantiFluor ONE dsDNA System (Promega, E4870) and Quantus Fluorometer (Promega, E6150).

Samples were diluted to 0.2 ng/μL, and Nextera XT DNA library preparation (Illumina, FC-131-1096) was carried out following the manufacturer’s instructions for a final volume of 25 μL. The DNA libraries were then normalized by the Heflin Genomics Core using qPCR and sequenced with a NovaSeq 6000. Sequencing reads were aligned to GRCz11 release 104 using the Lawson Lab Zebrafish Transcriptome Annotation version 4.3.2^34^ using the STAR aligner (2.7.3a-GCC-6.4.0-2.28). The resulting raw count files were normalized using rlog counts method in DESeq2. The code for downstream analysis, including differentially expressed genes, differentially expressed transcripts, and GO analysis, is available on Github: https://github.com/thymelab/znf804a_transcriptomics

### CRISPR/Cas9 Injection and RT-qPCR

Guide RNAs targeting the *znf804a* gene were generated using the MAXIscript T7 transcription kit (Thermo #AM1312). 100 ng of 3 guides targeting either regulatory or protein-coding elements of the *znf804a* gene were mixed, and a ratio of 1 uL of this guide RNA cocktail was added with 0.5 uL of Cas9 mRNA (at 1000 ng/uL), which was generated using the mMessage mMachine T7 Transcription Kit (Thermo #AM1344). Guide sequences and qPCR primers are listed in **Table S13**.

EK wild-type breeding fish were separated overnight and allowed to interact the following morning. Following 15-minutes, embryos were collected and injected with a cocktail of CRISPR guide RNA and Cas9 RNA as previously described^44^. At 6 dpf, 3 fish were pooled and RNA extraction was performed as described above. cDNA synthesis was performed using iScript Reverse Transcription Supermix for RT-qPCR (Bio-Rad #1708840) using 200 ng of RNA template. RT-qPCR was performed with SsoAdvanced Universal SYBR Green Supermix (Bio-Rad #1725271) with 4 ng of input using a Bio-Rad CFX96 Touch Real-Time PCR Detection System Module on a C1000 Touch Thermal Cycler. The *rpl13a* gene was used as the housekeeping gene. Gene expression was calculated using the comparative method delta-CT.

## Acknowledgments

We thank the Heflin Genomics Institute at UAB, Harvard, UAB, and UMass Chan fish facility staff, and the Research Computing teams at UAB and UMass Chan for supporting this study. We also thank members of the Thyme lab for technical support, and Jack Yuanwei Cheng for feedback on the manuscript.

## Author contributions

BLB, EHL, and SBT conceptualized the study. BLB, EHL, MESC, and SBT performed experiments. BLB and SBT performed data analysis. BLB and SBT wrote the original manuscript. All authors revised and approved the final version of the manuscript.

## Data and materials availability

All data are available in the main text, the Supplementary Materials, or appropriate databases. The *znf804a*^a364^ mutant is available from ZIRC. All code is available from https://github.com/thymelab. Behavioral data is available from Zenodo under DOI: 10.5281/zenodo.19701534. RNA-sequencing data, raw counts, and fastq files are available from GEO under the access number GSE253405.

