## Supplementary Figures 1-4 for "Znf804a is a regulator of circadian behaviors in zebrafish"

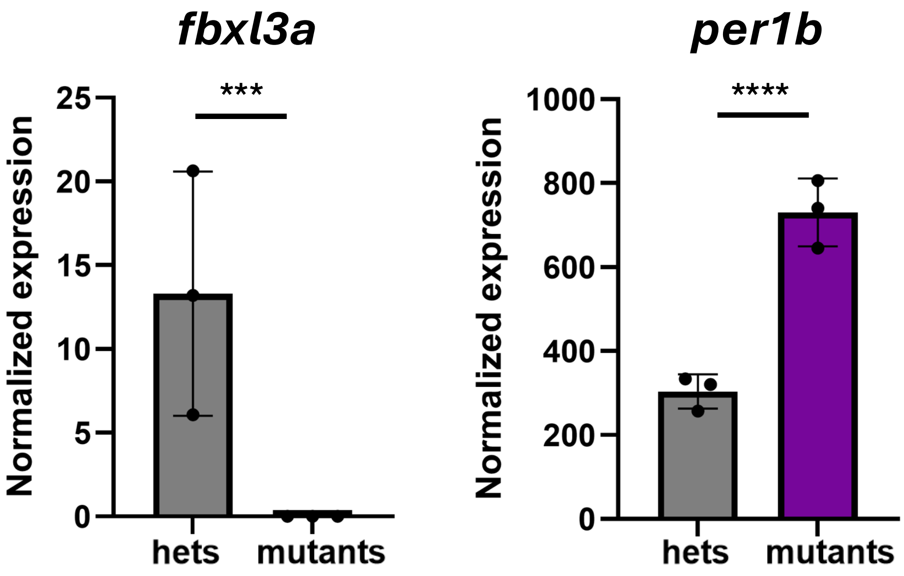


Figure S1: **Preliminary RNA sequencing data indicate disruptions in *fbxl3a***. Bar plot depicting heterozygous vs mutant normalized expression of *fbxl3a* and *per1b*. Significance is denoted by raw p-values from RNA seq analysis (*=p<0.05, **=p<0.01, ***=p<0.001, ****=p<0.0001)

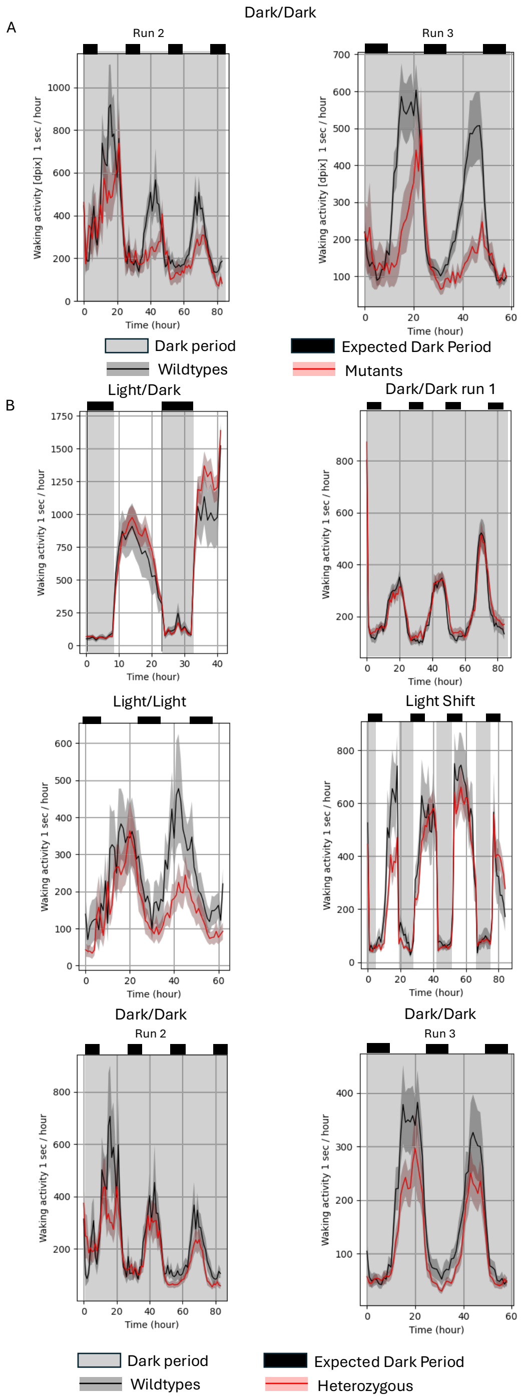


Figure S2: **Replication of circadian drift dark/dark conditions in *znf804a* mutants. A**) Wild-type (black) (n=21 run 2; n=32 run 2) and *znf804a* mutants (red) (n=19 run 2; n=16 run 3) waking activity binning by hour over the course of a replicate of the dark/dark experiment starting at 4 dpf. Like in the previous replicate (Figure 2B), the mutant zebrafish display drifts in circadian activity and reduced overall activity **B**) Wild type (black) and *znf804a* heterozygous (red) (n=40 light/dark; n=40 dark/dark run 1; n=34 light/light; n=30 light shift; n=41 dark/dark run 2; n=45 dark/dark run 3) waking activity for corresponding experiments.

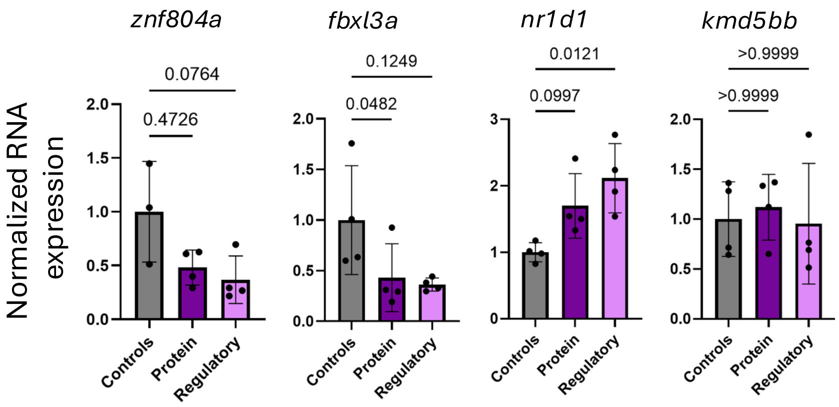


Figure S3: **CRISPR targeting of the protein-coding or regulatory region of *znf804a* influences circadian gene expression**. **A**) Targeting the regulatory region of *znf804a* leads to a trend-level ~65% reduction in *znf804a* RNA expression, while targeting the protein-coding region produced a 52% reduction that did not reach statistical significance, consistent with continued transcription despite coding disruption. **B**) Targeting the regulatory or protein-coding regions leads to consistent, though variable, reductions in *fbxl3a* expression. The variability is consistent with mosaicism in crispant embryos. **C**) Targeting of *znf804a* was also associated with increased *nr1d1* expression, consistent with our stable line's effect on *nr1d1*, supporting downstream effects on circadian gene expression. **D**) Expression of the control gene, *kmd5bb*, was unaffected, indicating the specificity of the perturbation. Samples were compared using a Kruskal-Wallis test corrected for multiple corrections against the control group.

Mutant v wildtype light/dark

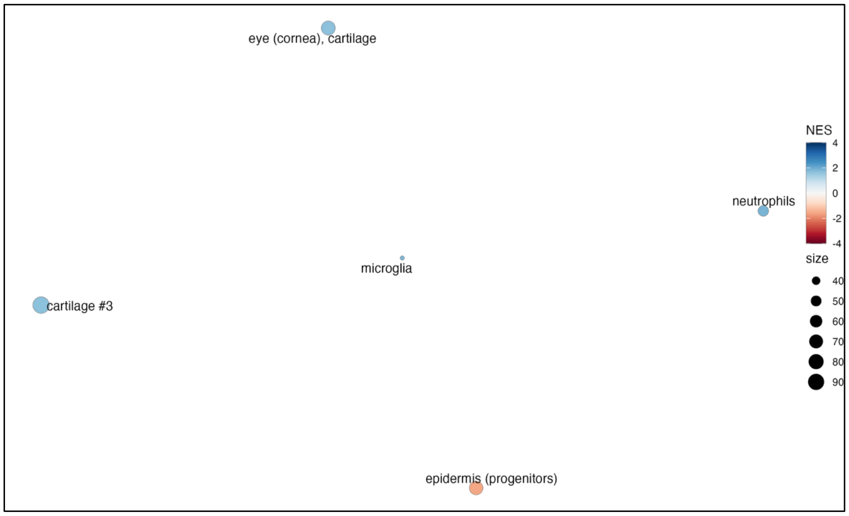


Mutant v wildtype dark/dark

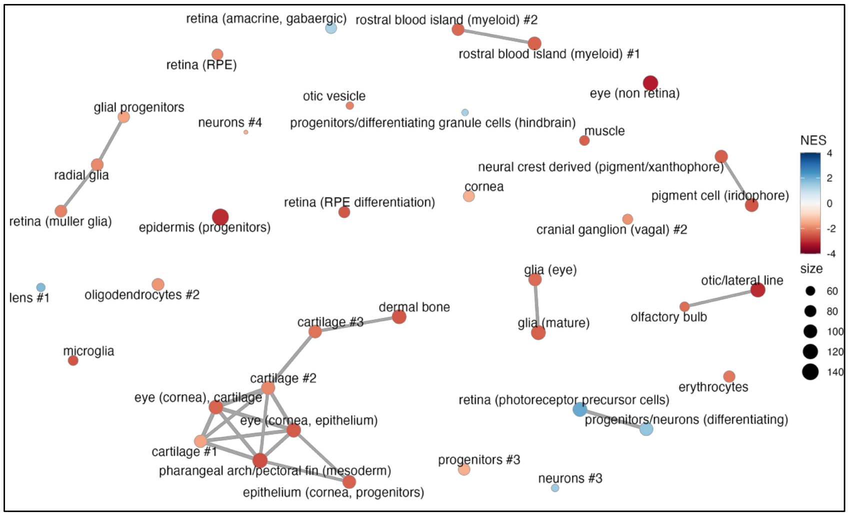


Figure S4: **Gene set enrichment analysis for mutant versus wildtype**. Fish were raised under normal light/dark (top) and dark/dark (bottom) conditions. Each node represents a cell-type specific gene set, the color represents the normalized enrichment scores (NES), and the size of the node represents the number of genes in the gene set.

Supplementary Tables

Table S1: **DEG for mutants compared to wildtypes raised under normal light/dark conditions.**

Table S2: **DEG for mutants compared to wildtypes raised under dark/dark conditions.**

Table S3: **DEG for wildtypes raised in dark/dark conditions compared to light/dark conditions**

Table S4: **DEG for mutants raised in dark/dark conditions compared to light/dark conditions**

Table S5: **Full table of Gene Ontology Analysis Terms for All 4 Group Comparisons**

Table S6: **Table of pairwise comparisons for circadian genes extracted from DEG comparisons of RNA seq**

Table S7: **DET for mutants and wildtypes raised under normal light/dark conditions**

Table S8: **Table of Gene Set Enrichment Analysis findings**

Table S9: **DEG for heterozygous fish compared to wildtypes raised under normal light/dark conditions.**

Table S10: **DEG for mutants compared to heterozygous fish raised under normal light/dark conditions.**

Table S11: **DEG for heterozygous fish compared to wildtypes raised under dark/dark conditions.**

Table S12: **DEG for mutants compared to heterozygous fish raised under dark/dark conditions.**

Table S13: **Table of Guide RNAs used for crispant injection**
